# Structural and functional analysis of TREM2 interactions with amyloid beta reveal molecular mechanisms that mediate phagocytosis of oligomeric amyloid beta

**DOI:** 10.1101/2024.05.23.595586

**Authors:** Jessica A. Greven, Omar Osario, Jay C. Nix, Jennifer M. Alexander-Brett, Tom J. Brett

## Abstract

The TREM2 receptor is expressed on microglia in the brain, where it plays critical roles regulating microglia function. TREM2 engages a number of ligands involved in Alzheimer’s disease, and consequent signaling triggers phagocytosis, activation, survival, and proliferation. TREM2 has emerged as a drug target for AD, however very little is known regarding the structural basis for TREM2 microglial functions. Here we investigated the engagement of oligomeric amyloid beta (oAβ42) with TREM2. Using familial variants of amyloid beta, we show that mutations in the N-terminal portion of Aβ, notably residues H6 and D7, disrupt binding to TREM2. We then co-crystallized TREM2 with Aβ(1-8) peptide and determined the high resolution crystal structure. The structure revealed the peptide binds to the hydrophobic site of TREM2, closest to CDR1. Mutational and binding studies using BLI confirmed that mutations to the hydrophobic site ablate binding to oAβ42. Finally, we show that these interactions are critical to triggering phagocytosis of oAβ42, as oAβ42 variants H6R and D7N are not phagocytosed. Altogether, these data indicate that TREM2 engages oAβ42 using the hydrophobic site on TREM2 and the N-terminal portion of Aβ, and that this interaction is critical to trigger signaling and phagocytosis.

## Introduction

Alzheimer’s disease (AD) is hallmarked by accumulation of extracellular amyloid beta (Aβ) plaques and deposition of intracellular hyperphosphorylated tau in neurofibrilary tangles (NFT)[1-4]. During disease progression, brain resident microglia cluster around extracellular Aβ plaques to inhibit further growth and deposition, thus limiting neuronal toxicity [5]. The TREM2 receptor expressed on microglia appears to play a critical role in these functions, and TREM2 has been shown to bind oligomeric Aβ (oAβ) and mediate phagocytosis of oAβ [6-8]. TREM2 knockouts in the 5XFAD AD mouse model further support a critical role for TREM2 managing Aβ plaques. In these models, microglia do not surround Aβ deposits, and these deposits are less dense, more diffuse and thought to be made of increased amounts of the more neurotoxic oligomeric Aβ [9]. Thus TREM2 expressed on microglia appear to play central roles in directing microglia functions to manage amyloid beta aggregates.

The Aβ42 peptide is created from proteolytic processing of the amyloid precursor protein (APP). Missense mutations in APP can result in Ab42 variants that are associated with early onset AD, or cerebral amyloid angiopathy (CAA), where amyloid deposits in cerebral vessels and can cause hemorrhagic stroke. Various point mutations in Aβ42 have been documented, including the English (H6R)[10], Japanese (D7N)[11], Flemish (A21G)[12], Dutch (E22Q)[13], Arctic (E22G)[14], Italian (E22K)[15], and Iowa (D23N)[16] mutations. These variants are usually observed to enhance Aβ aggregation kinetics [17]. It is unknown whether these variants impact interactions with TREM2 or TREM2-mediated microglial functions.

Despite the importance that TREM2 interactions with Aβ appear to have on microglial function and AD pathology, little is known regarding the molecular details that govern this interaction. Here, we use comprehensive biophysical and structural methods to characterize this interaction, and identify the TREM2 surface and Aβ peptide region that is most critical in mediating binding. Finally, we show that these interactions are critical to mediating microglia phagocytosis of oligomeric Aβ42 (oAβ42). Altogether, these results not only delineate critical determinants of interaction that limit AD pathology, but they also provide a framework for therapeutically targeting TREM2 in limiting AD progression.

## Methods

### Protein expression and purification

Throughout this work, the following TREM2 constructs are defined: TREM2 = a.a. 19-134; sTREM2 = 19-157; MBP-TREM2 = MBP fused to TREM2 19-134 as described in [18]; TREM2 receptor = 1-230. MBP-TREM2 N79Q was made by Q5 site directed mutagenesis. TREM2 and sTREM2 variants were synthesized as gBlocks and cloned into pHLsec vector[19] by Gibson Assembly. All TREM2 proteins were produced as secreted proteins from Expi293 cells and purified from media as previously described [20]. Biotinylated TREM2 proteins were purified by NiNTA chromatography and enzymatically biotinylated using BirA enzyme as described previously [21, 22]. Biotinylated TREM2 proteins were immediately aliquoted, frozen, and stored at -80 ºC. sTREM2 proteins were purified by NiNTA chromatography, followed by gel filtration using an s200 increase column and running buffer consisting of 20mM Tris pH 8.5, 150mM sodium chloride, and 0.01% sodium azide. Biotinylated TREM2 proteins and sTREM2 proteins were immediately aliquoted, frozen, and stored at -80 ºC. For reproducibility, freshly thawed aliquots were used for each experiment. MBP-TREM2 N79Q was purified by NiNTA chromatography, followed by gel filtration using an s200 increase column and running buffer consisting of 50mM HEPES pH 7.5, 250mM sodium chloride, and 0.01% sodium azide.

### Production of WT and familial variant oAβ42 oligomers

WT Aβ42 as well as English (H6R), Japan (D7N), Dutch (E22Q), Arctic (E22G), Flemish (A21G), Iowa (D23N), and Italian (E22K) variant peptides were purchased from Anaspec. Oligomers were produced similar to previous descriptions [23]. Briefly, peptides were first dissolved in hexafluoroisopropanol (HFIP) and allowed to air-dry overnight. They were then dissolved in DMSO at a concentration of 100 mM and sonicated for 10 min. These solutions were then diluted into PBS at a concentration of 22 µM and incubated at 37 ºC for 3 hours. Precise concentrations were then measured by BCA assay, after which the oAβ42 solutions were aliquoted, flash frozen, and stored at -80 C. Freshly thawed aliquots were used for each experiment. Monomeric biotinylated oAβ42 was purchased from Anaspec and then mixed 1:20 with non-biotinylated monomeric Aβ42. These monomers were then oligomerized using the previously stated protocol.

### Transmission Electron microscopy

oAβ42 preparations were characterized by negative stain transmission electron microscopy (TEM). Samples were prepared as previously mentioned and diluted to final Aβ42 concentrations of 2.5 μM. These TEM samples (10 μl) were applied to carbonate-coated grids for 1 minute and negatively stained with 1% phosphotungstic acid (PTA) for 1 minute. TEM micrographs were obtained on a Hitachi H-7000 operated at 120 kV.

### Binding studies by biolayer interferometry (BLI)

BLI data were collected on an Octet RED384 system (FortéBio). Biotinylated oAβ42 and TREM2 bound to ForteBio Streptavidin-coated biosensors and binding kinetics were measured using a running buffer of PBS with 0.1% BSA and 0.005% Tween-20. Data were processed using double-reference subtraction (loaded protein into buffer and biotin-loaded pin into ligand) in ForteBio Data Analysis 9.0.

### MBP-TREM2 N79Q crystallization, structure determination, and analysis

MBP-TREM2 N79Q was concentrated to 15 mg/mL. Crystals were grown over 3-5 days at 17 ºC by hanging drop vapor diffusion by mixing 1:1 with a well solution containing 11% PEG 4000 and 100 mM sodium citrate at pH 5.5 and streak seeding. Amyloid beta peptide (1-8: DAEFRHDS) from GenScript was co-crystallized with MBP-TREM2 N79Q by mixing at an 18:1 molar ratio prior to crystallization. Crystals were cryoprotected in 15% PEG 4000, 20% PEG 400, and sodium citrate at pH 5.5 and flash frozen under a liquid nitrogen stream at -160 ºC. Data were collected at the Advanced Light Source beamline (Berkeley, CA) and processed to 1.84 Å resolution using XDS[24]. Data collection and refinement statistics are given in Table 1. Crystals had similar unit cell and space group parameters to the previously published MBP-TREM2 structure [18], so the phase problem was solved by isostructural replacement using protein coordinates from PDB ID 6XDS. The initial solution was refined by rigid body refinement in PHENIX[25] and the model was improved in subsequent rounds of refinement conducted with individual B factors and iterative rebuilding through COOT [26]. Secondary structure restraints were used during refinement and in later rounds of refinement, optimization of X-ray and ADP or stereochemistry weight was applied. LigPlot+ [27] was used to analyze the TREM2 - Aβ interface. All crystallographic and analysis software were compiled through the SBGrid resource [28] and diffraction images were archived with the SBGrid Data Bank [29]. The coordinates and experimental structure factors have been deposited in the RCSB Protein Data Bank.

**Table 1.** Crystallographic statistics for human MBP TREM2 N79Q/Aβ(1-8) complex.

| **Table 1. Crystallographic statistics for human MBP TREM2 N79Q/Aβ(1-8) complex** |  |
| --- | --- |
| **Data collection statistics** |  |
| Space Group | P 2<sub>1</sub> 2<sub>1</sub> 2<sub>1</sub> |
| Unit Cell (Å and °) | a=73.63 a = 90<br>b=79.46 b = 90<br>c=103.68 g = 90 |
| Source | ALS 4.2.2 |
| Wavelength(Å) | 1.072 |
| Resolution(Å) | 47.90 – 1.84 (1.88-1.84) |
| R<sub>merge</sub> | 0.129 (2.372) |
| CC<sub>1/2</sub> | 0.997 (0.394) |
| Completeness(%) | 100 (100) |
| Redundancy | 7.0 (6.9) |
| I/$\sigma$(I) | 8.7(0.7) |
| **Refinement statistics** |  |
| R<sub>work</sub> (%) | 19.42 |
| R<sub>free</sub> (%) | 22..61 |
| Amino Acid Residues(#) | 497 |
| Waters (#) | 308 |
| RMSD bond length (Å)/angles(°) | 0.005/0.784 |
| Wilson B (Å<sup>2</sup>) | 31.25 |
| Ave. B protein (Å<sup>2</sup>) | 39.59 |
| Ave. B water (Å<sup>2</sup>) | 42.80 |
| **Ramachandran** |  |
| %Favored | 97.55 |
| %Allowed | 2.04 |
| %Outliers | 0.41 |
| Rotamer outliers (%) | 3.12 |
| Clashscore (score / percentile) | 3.80 |

### HMC3 phagocytosis assay

Frozen human microglial cells (HMC3) from American Type Culture Center were thawed and grown to confluency in EMEM media containing 10% FBS and 1% penicillin/streptomycin at 37 ºC and 5% CO_2_ . Well slides were coated with fibronectin/collagen and incubated for 2 hours to enhance cell adhesion. Cells were then seeded into wells at 80,000 cells/well. As a control, half of the wells were pretreated for 1 hour at 37 ºC with Cytochalasin D in DMSO at a concentration of 10uM with 0.01% DMSO in solution to block actin polymerization for phagocytosis. The other half of the wells were incubated with 0.01% DMSO for 1 hour at 37 ºC. After pretreatment, cells were incubated for 2 hours at 37 ºC with oAβ42 WT, H6R, D7N, and A21G at a concentration of 1 μM in EMEM media. After incubation, cells were washed with PBS and fixed in 4% paraformaldehyde for 10 minutes and stored overnight at 4 ºC. Cells were then permeabilized in Tris buffered saline with 0.1% Triton (TBST) for 10 minutes. Coverslips were then blocked for 1 hour in a blocking solution (PBS with 2% BSA and 2% fish gel) and then incubated with anti-amyloid beta primary antibody (DE2B4) from Abcam diluted 1/400 in the blocking solution. Coverslips were subsequently washed in TBST for 10 minutes and then amplified using VectaFluor anti-mouse IgG amplification kit according to protocol. Lastly, coverslips were mounted on a microscopy slide using VECTASHIELD mounting medium for fluorescence with DAPI (Vector lab, #H-1200). Fixed cell imaging was performed on an Olympus 1X83 inverted microscope with 40 and 60X objectives and a Hammamatsu Orca camera.

## Results

### TREM2 binding to oAβ42 is severely inhibited for oAβ42 familial N-terminal variants

In order to investigate whether familial variants in oAβ42 impacted interaction with TREM2, we carried out quantitative binding studies using BLI. For these studies, biotinylated TREM2 proteins were immobilized on streptavidin biosensors and probed for binding to oAβ42 WT and variants in solution (**Fig 1A,B**). We found that WT oAβ42 bound TREM2 with very high affinity (K_D_ = 42.5 ± 0.6 nM) (**Fig. 1C**), similar to what has been reported previously using similar experimental orientations [30]. We next probed for binding to oAβ42 containing reported familial variants. These variants can be roughly grouped as occurring mid-peptide (Dutch E22Q, Arctic E22G, Flemish A21G, Iowa D23N, and Italian E22K) or in the N-terminal portion of the peptide (English H6R and Japan D7N). For mid-peptide variants, we found that most of these variants (E22Q, E22G, A21G, D23N) bound with similar binding magnitude to WT (**Fig 1D-G**). The exception was Italian E22K, which did not display any binding at the concentrations probed (**Fig 1H**). We next probed TREM2 binding to the N-terminal variants. Surprisingly, the N-terminal variants displayed very little (H6R) or no (D7N) binding (**Fig.1I,J**). Transmission electron microscopy (TEM) was used to validate that all WT and variant samples contained oligomers (Supplemental Figure 1). These results show that TREM2 binds WT oAβ42 with high affinity, and suggest that the interaction is mediated mainly by determinants in the N-terminal portion of the Aβ42 peptide.

**Figure 1.**
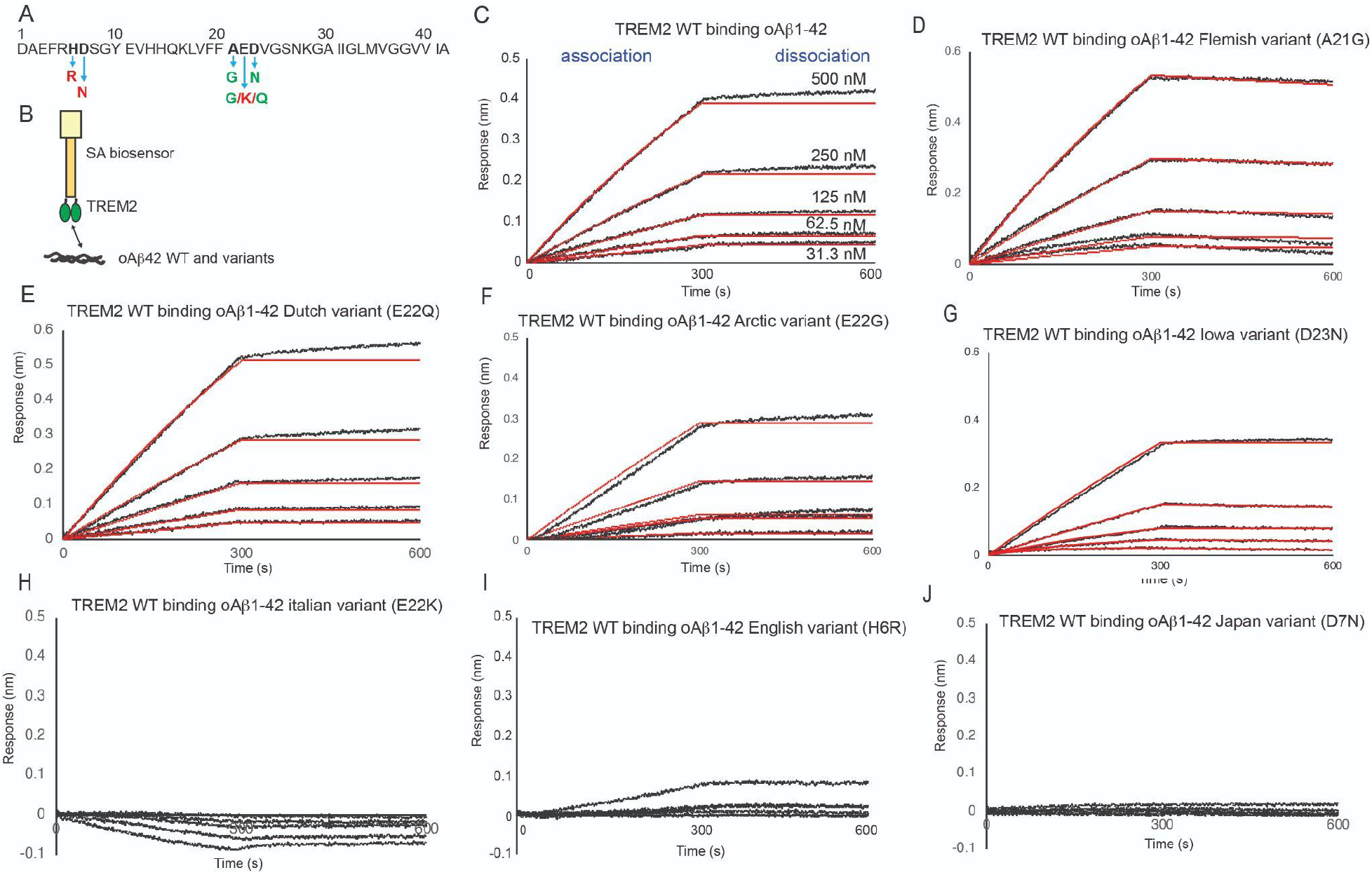
Familial mutations in N-terminal region of oligomeric Aβ42 alter binding to TREM2. **(A)** Sequence of Ab42 peptide showing position and sequence of familial variants. Point variants are colored as: red: ablated binding; green: did not impact binding. **(B)** Scheme of experiment. Immobilized TREM2 was probed for binding to oligomeric Aβ42 and familial variants (500 – 31.25 nM).**(C)** BLI response for TREM2 WT binding to oAβ42 or **(D)** Flemish variant (A21G), **(E)** oAb42 Dutch variant (E22Q), **(F)** Arctic variant (E22G), **(G)** Iowa variant (D23N), **(H)** Italian variant (E22K) **(I)** English variant (H6R), and **(J)** Japan variant (D7N) .

### Structure of TREM2 in complex with Aβ1-8

We then used the results of the BLI studies with oAb42 familial variants to design a minimal Aβ peptide that could be co-crystallized with TREM2, in order to structurally characterize this interaction. Since our binding results indicated that TREM2 interactions were critically mediated by the N-terminal portion of the amyloid beta peptide, we utilized the first 8 amino acids, Aβ(1-8), in co-crystallization experiments with an MBP-TREM2 fusion protein that had previously been demonstrated to produce crystals that diffract to high resolution [18]. We were able to obtain crystals of the complex that diffracted to 1.84 Å (Table 1). Difference electron density corresponding to the Aβ(1-8) peptide was observed on the distal end of TREM2 near the CDR1 loop (**Fig. 2A**). Our BLI studies with oAβ42 variants indicated critical importance of residues H6 and D7. Aβ D7 was observed to make pi-stacking interactions with TREM2 W44 and van der Waals contacts with TREM2 M41 in CDR1 (Fig. 2B and Supp. Fig. 2). It is possible that the D7N mutation might drive repulsion of W44 due to the charge change from Asp to Asn. Aβ H6 engaged in van der Waals contacts with TREM2 K42. The H6R variant would likely drive repulsion from K42. These observations suggest why the H6R and D7N oAβ42 variants might not engage TREM2. Further, the structure revealed hydrogen bond contacts for TREM2 R46 with multiple backbone residues in the Aβ peptide. A majority of the rest of the contacts are van der Waals interactions with residues in CDR1 as well as some in the F-G loop. The CDR2 is partially disordered and lies adjacent to the CDR1 loop. Overall, the crystal structure suggests the basis for the oAβ42 H6R and D7N variants to show disrupted binding to TREM2, and reveals critical contacts in CDR1.

**Figure 2.**
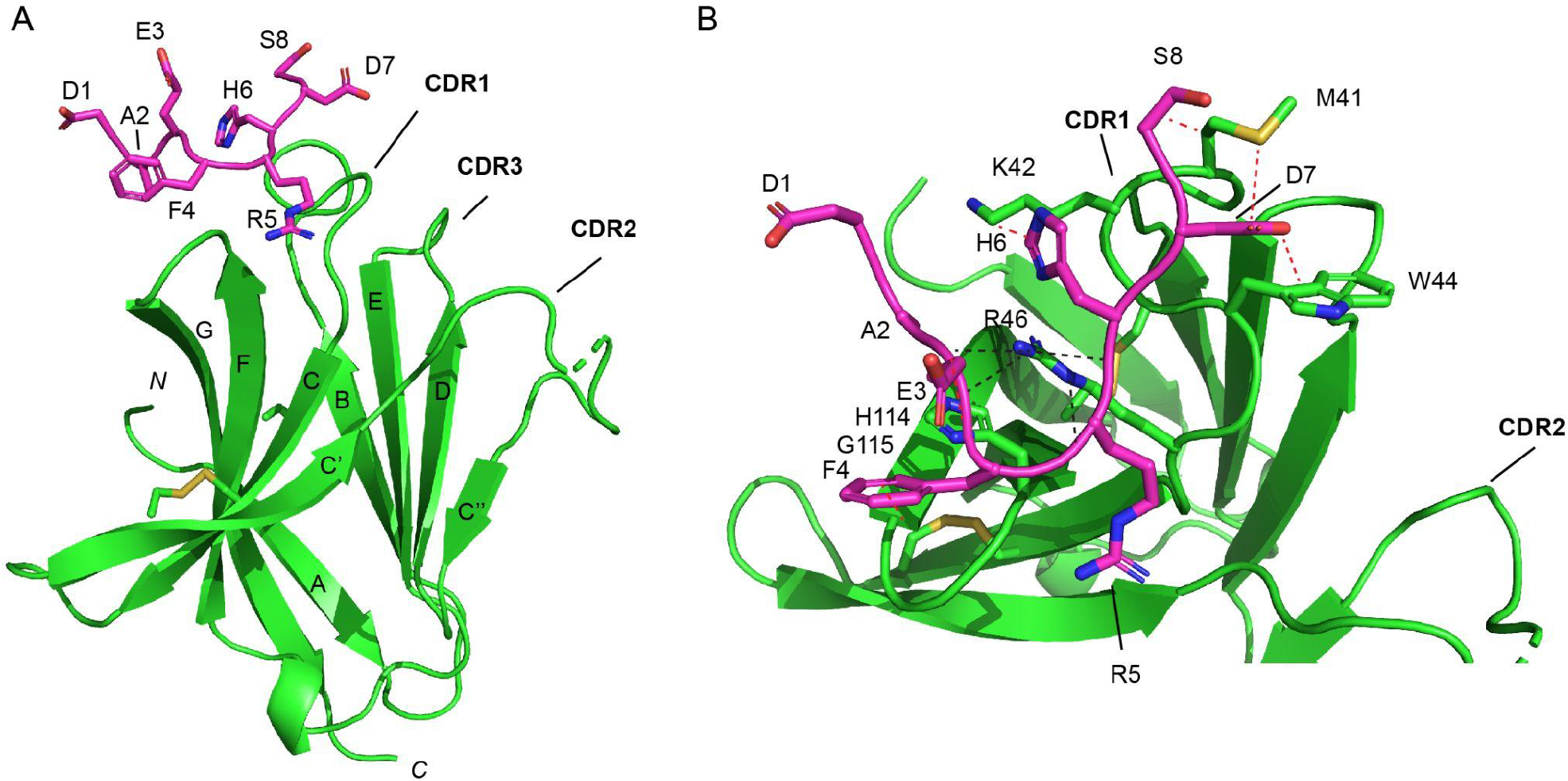
Crystal structure of Aβ(1-8) in complex with MBP-TREM2. **(A)** Overall structure of TREM2 (green) with Aβ(1-8) (magenta). MBP and the C-terminal 6His tag has been omitted for clarity. Aβ1-8 peptide binds near CDR1 loop, adjacent to CDR2 loop. **(B)** View showing binding site on TREM2 for Aβ(1-8). Van der Waals contacts between residues are denoted with red dashes while hydrogen bonds are denoted by black dashes.

### BLI validation of binding site

In order to validate our findings of the TREM2-Aβ1-8 crystal structure, we performed biophysical binding experiments using BLI and variant TREM2 proteins. Since we observed TREM2 engaging the Aβ1-8 peptide in the hydrophobic site, we first analyzed variants in this region. We analyzed point mutations in CDR1 (M41D), CDR2 (L69D, W70D, and L71D) and CDR3 (L89D). We found M41D displayed reduced binding, consistent with M41 being a contact residue with Aβ(1-8) peptide in the crystal structure (**Fig 3B,C**). We also found that point variants in the CDR2 loop - L69D, W70D, and L71D - all displayed notably decreased binding to oAβ42 compared to TREM2 WT (**Fig 3B-G**). Since CDR2 is nestled adjacent to CDR1, and CDR2 displays conformational flexibility and disorder, we reasoned that these mutations might influence CDR1 conformations. To further probe this, we created a double mutant in the CDR2 loop (L69D/L71D) and triple mutant targeting the CDR1 and CDR2 loops (W44D/L69D/L71D). Remarkably, we found that these variants displayed little to no binding to oAb42 at the concentrations probed (**Fig. 3 H,I)**. The triple mutant was the most disruptive, which reflects the critical role of W44 in the crystal structure of the complex. Mutation to Asp (W44D) would create a charge repulsion with D7. For comparison, we investigated oAβ42 binding to basic site point variants T96K, R47H. We found that R47H displayed only slightly reduced binding (**Fig. 3L)**, while binding to TREM2 T96K was more reduced (**Fig. 3K)**. Due to the importance of R46 in the crystal structure of the complex, we probed binding to the double mutant R46A/R47A and found it still retained binding to oAβ42. These results validate that oAβ42 primarily engages the hydrophobic site on TREM2 as elucidated in our co-crystal structure.

**Figure 3.**
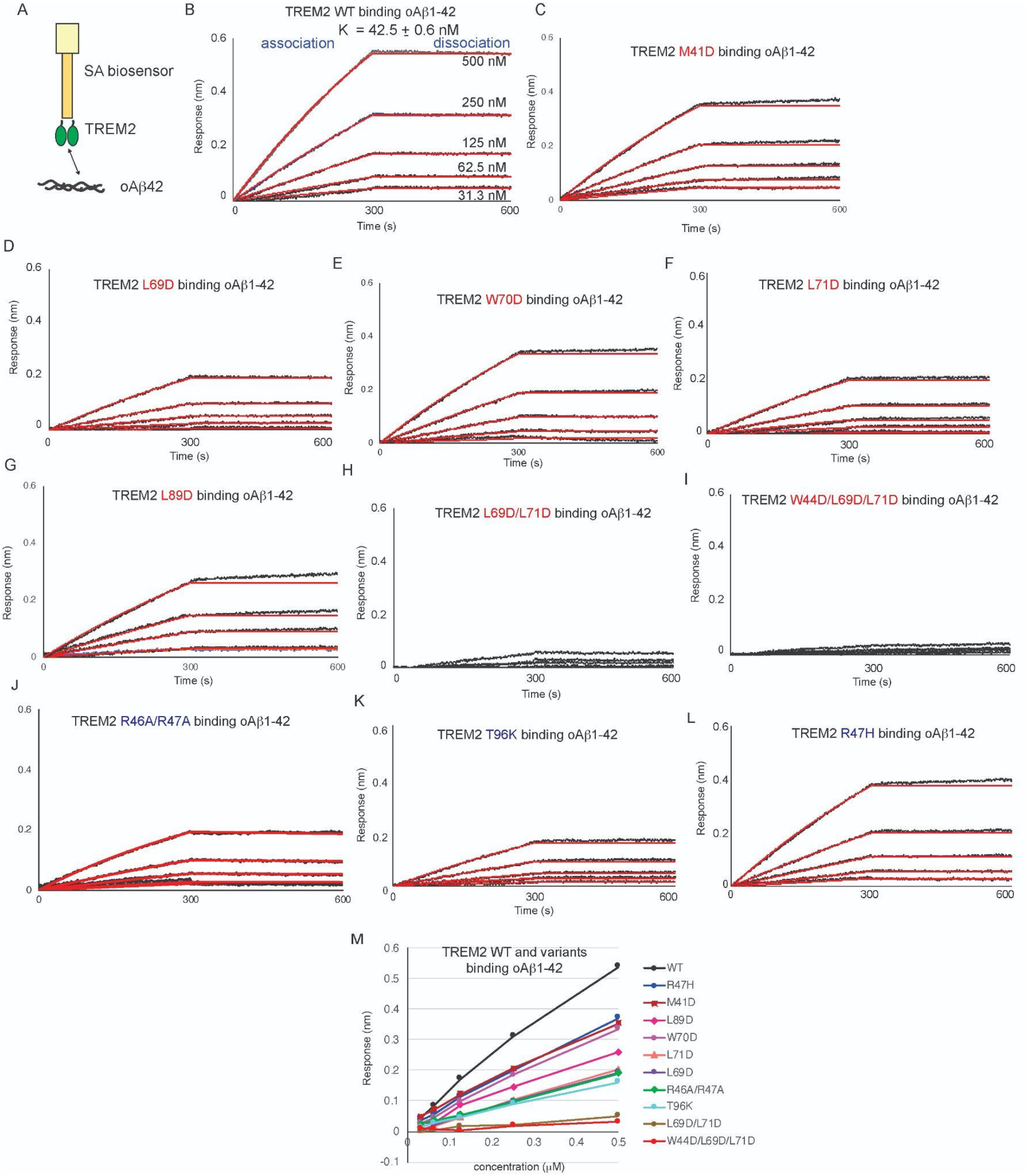
Mutations to TREM2 hydrophobic site disrupt binding to oligomeric Ab42. Immobilized TREM2 WT and variants were probed for binding to oligomeric Ab42 (500 – 31.25 nM). (A) Scheme of experiment. **B-L)** BLI response for TREM2 **B)** WT, **C)** M41D, **D)** L68D, **E)**, W70D, **F)** L71D, **G)** L89D, **H)** L69D/L71D, **I)** W44D/L69D/L71D, **J)** R46A/R47A, **K)** T96K, **G)** R47H binding to oAb42 (31.3 - 500 nM). Double-reference subtracted data (black) overlayed with 1:1 kinetic fits (red). K_D_ derived from kinetic fits. Little to no binding is detected for L69D/L71D and W44D/ L69D/L71D. Data representative of at least two independent experiments. **M)** BLI Max response plotted versus concentration for TREM2 WT and all variants.

Since the orientation in our BLI experiments would allow for a single oAβ42 molecule to engage multiple immobilized TREM2 molecules on the pin, we reasoned that switching the orientation of the experiment (by immobilizing oAβ42) would allow for more dramatic differences in binding. Thus, we carried out experiments with immobilized oAβ42 binding to sTREM2 WT and variants (**Fig. 4A**). In this orientation, point mutations to the hydrophobic site (L69D, L71D) as well as double mutations (L69D/L71D) displayed no binding to oAβ42 (**Fig. 4F-H)**. In contrast, mutations to the basic site displayed mild (R47H, R62H) or little (R46A, R46D) reduction in binding (**Fig 4D-E, I-J)** compared to WT (**Fig 4C)**. Altogether, these results demonstrate that oAβ42 primarily engages the TREM2 hydrophobic site.

**Figure 4.**
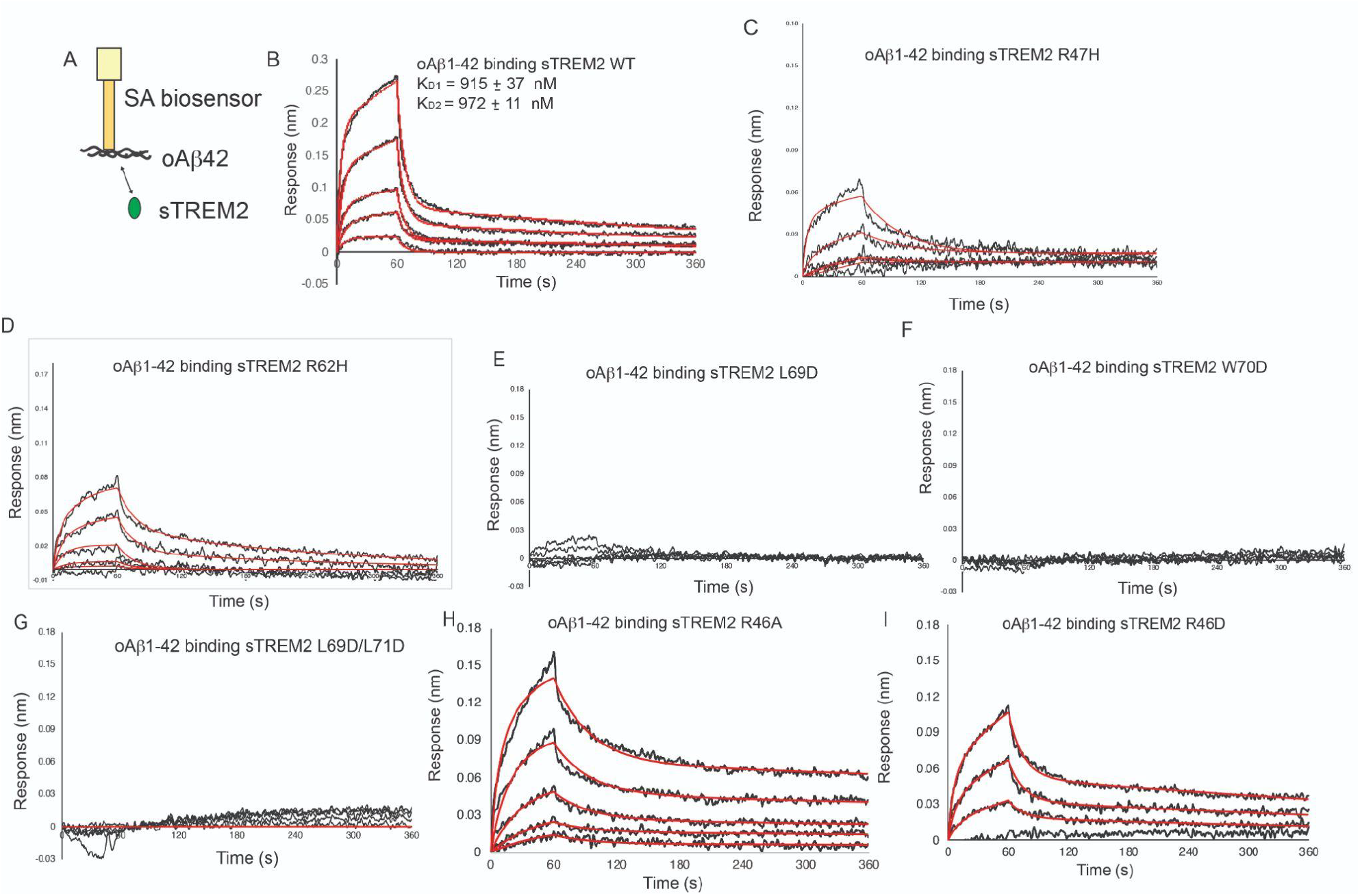
Mutations to the hydrophobic site prevent sTREM2 binding to oAβ42, mutations to the basic site do not. **A)** Scheme of experiment. **B-I)** BLI response for sTREM2 **B)** WT, **C)** R47H, **D)** R62H, **E)** L69D, **F)** W70D, **G)** L69D/L71D, **H)** R46A, and **J)** R46D. binding to oAβ42. sTREM2 concentration range was 0.0625 - 1.0 μM. Double-reference subtracted data (black) overlayed with 2:1 kinetic fits (red). Data representative of at least two independent experiments.

### Structural determinants of phagocytosis

To characterize functional ramifications of oAβ42 interactions with TREM2, we carried out phagocytosis assays using the HMC3 human microglial cell line. In order to address the importance of N-terminal Aβ42 interactions with TREM2 in mediating phagocytosis, we utilized oligomers containing N-terminal familial variants H6R and D7N. For comparison, we utilized oAβ42 WT and the A21G variant. HMC3 cells were treated with 1 uM oAβ42 WT and variants, incubated for 2 hours at 37C, and then assessed for phagocytosis by immunofluorescence using the anti-Aβ42 antibody DE2B4, whose epitope is in residues 8-16, and therefore should recognize all variants studied here. Cytochalasin D (CytoD), a phagocytosis inhibitor, was used to indicate involvement of phagocytosis in internalization. Significant amounts of oAβ42 WT uptake were detected, and these were largely blocked by CytoD treatment (Fig. 5). Uptake can be seen in the WT Aβ experiment. In comparison, moderate levels of oAβ42 A21G internalization were observed. In stark contrast, almost no uptake of oAβ42 H6R or D7N were observed, consistent with our BLI binding studies which indicated little to no TREM2 binding for these variants. These results indicate that TREM2 interactions with the N-terminal portion of Aβ are critical to driving phagocytosis in microglia.

**Figure 5.**
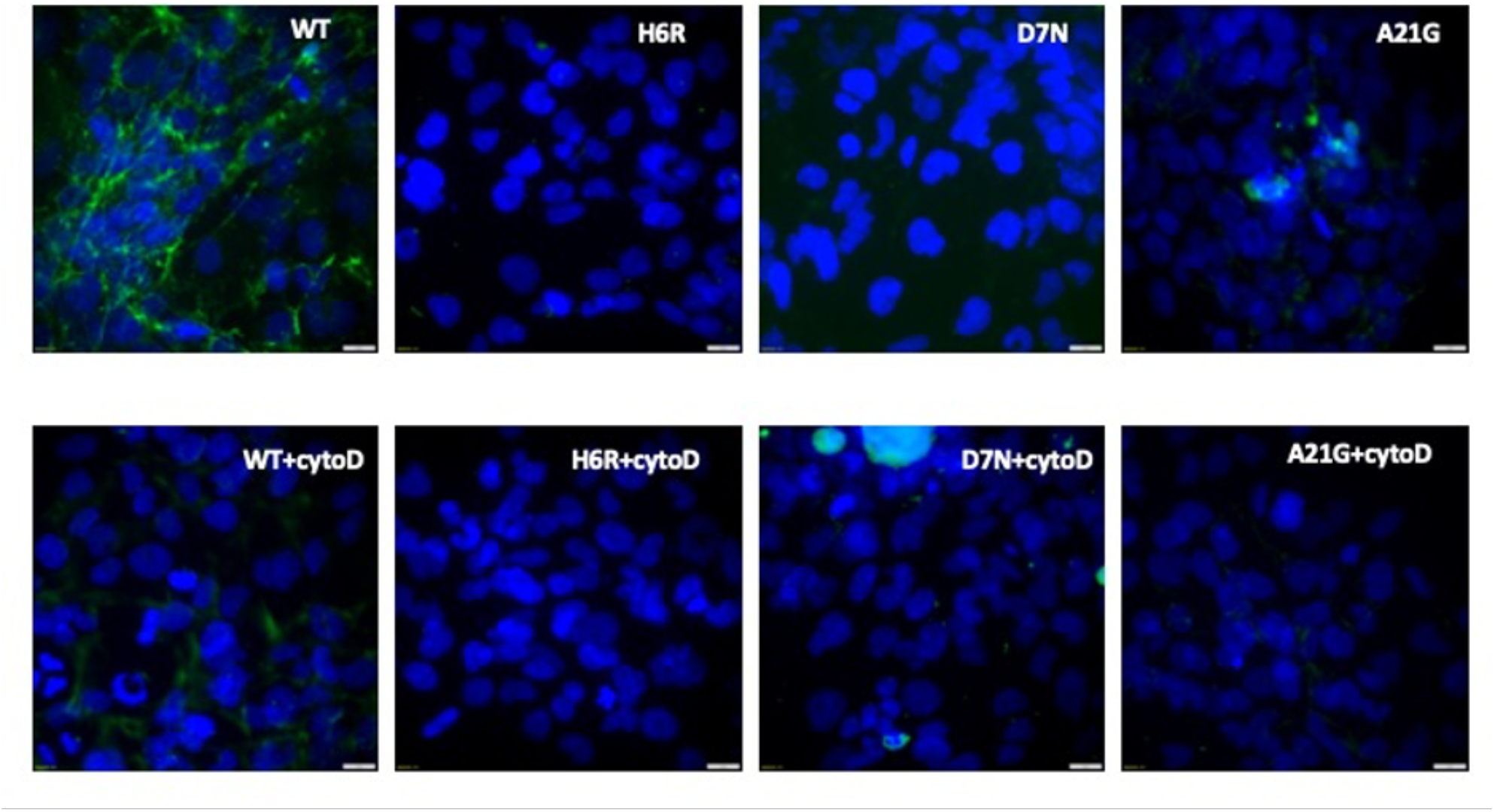
Involvement of N-terminal Ab42 residues in mediating microglia phagocytosis of oAb42. Representative immunocytochemical staining images of HMC3 cells after pre-treatment with vehicle (DMSO) (top row) or cytoD treatment for 2 hours followed by treatment with 1 uM oAb42 WT or the H6R, D7N, or A21G familial variants. Cells were stained with DAPI (blue-nuclei) and Ab antibody DE2B4 (green).

## Discussion

To our knowledge, this is the first study revealing the structural mechanisms of Aβ interaction with a phagocytic microglial receptor. However, there have been limited structural studies of Aβ interactions with neuronal receptors. In microglia, binding of Aβ to TREM2 induces downstream signaling for microglia activation and phagocytosis which are beneficial to remove oAβ and regulate Aβ plaque formation [9]. Thus one would want to therapeutically target this interaction with mimics or enhancers. In contrast, oAβ engagement of other neuronal receptors triggers neurotoxic signaling and there have been efforts to understand these interactions in order to develop inhibitors [31]. Around 10 different Aβ neuronal receptors have been identified to date, with PrPc, FcgammaRIIb, and LilrB2 being the most well-studied. A structural study of LilrB2 illustrates that the receptor engages the middle portion of the Aβ peptide (16 KLVFFA 21) [32]. A high-resolution spectroscopic study of PrPc, FcgammaRIIb, and LilrB2 showed that these receptors all bind to the growing end of Aβ fibrils, though specific residues were not identified[33].

In contrast, our results suggest that TREM2 engages the N-terminal portion of the Aβ peptide. Binding data indicates that point mutations at the N terminal portion of Aβ ablate binding, while mutations at mid-peptide residues typically have no impact on binding. These results corroborate our crystal structure where the hydrophobic site of TREM2 makes contact with oAβ at its N terminus. The functional implications of this finding were probed using a phagocytosis assay to determine how N terminal mutations on Aβ impacted uptake into HMC3 microglial cells. Results from the assay displayed minimal uptake at oAβ42 N terminal variants while mid peptide mutation displayed some, albeit reduced, uptake to WT. Because the only published studies investigating oAβ42 engagement with receptors report binding at mid peptide regions, these results are novel and incredibly impactful to understanding how Aβ engages a variety of receptors through unique determinants throughout the peptide.

Our structural and mutational studies indicate that TREM2 utilizes the hydrophobic site to engage oAβ42. While our structural studies revealed that the major contacts for the Aβ(1-8) peptide were in CDR1, we also found that mutations to the CDR2 loop caused the most dramatic decrease in binding to oAβ42. These observations suggest that the conformationally dynamic CDR2 likely influences CDR1 through direct contacts, thus influencing the ability of CDR1 to attain conformations optimal for binding oAβ42. They also indicate that targeting the CDR2 loop with small molecule drugs might represent a way to modulate the binding of TREM2 to oAβ42. Thus, not only do our studies define how TREM2 engages oAβ42 to trigger phagocytosis, they also provide a structural and conceptual framework for targeting TREM2 functions with small molecules.

## Acknowledgements

This work was supported by Alzheimer’s Association Research Grant (AARG-16-441560)(TJB), Bright Focus Foundation (A2022032S)(TJB), and Amgen Scholars Research Fellowships (JAG).

---

**Figure 15.**
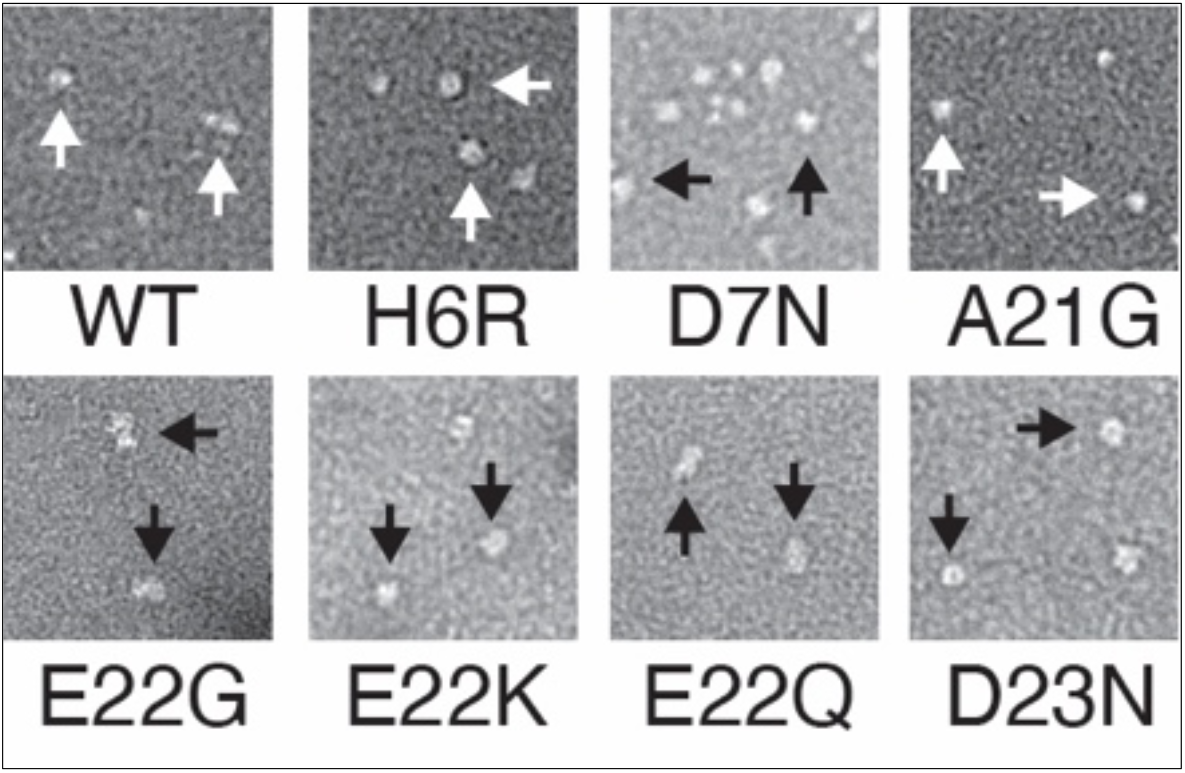
TEM of preps of variant oAβ42. Images shown at 30,000X magnification. Arrows denote some of the amorphous oligomers seen. These preps were used in BLI binding studies

